# Preventive training interferes with mRNA-encoding myosin 7 and collagen I expression during pulmonary arterial hypertension

**DOI:** 10.1101/2020.12.17.423207

**Authors:** Thaoan Bruno Mariano, Anthony César de Souza Castilho, Ana Karenina Dias de Almeida Sabela, André Casanova de Oliveira, Sarah Santiloni Cury, Andreo Fernando Aguiar, Raisa de Jesus Dutra Dias, Antonio Carlos Cicogna, Katashi Okoshi, Luis Antonio Justulin Junior, Robson Francisco Carvalho, Francis Lopes Pacagnelli

## Abstract

To gain insight on the impact of preventive exercise during pulmonary arterial hypertension (PAH), we evaluated the gene expression of myosins and gene-encoding proteins associated with the extracellular matrix remodeling of right hypertrophied ventricles. We used 32 male Wistar rats, separated in four groups: Sedentary Control (S; n=8); Control with Training (T; n=8); Sedentary with Pulmonary Arterial Hypertension (SPAH; n=8); and Pulmonary Arterial Hypertension with Training (TPAH; n=8). The rats trained for thirteen weeks on a treadmill. They had two weeks of adaptation training. The PAH was induced by application of monocrotaline 60 mg/kg. Consequential right ventricular dysfunction was observed after the 10th week of training. Rats in the control group received saline application. At the end of the 13th week, echocardiography analysis confirmed cardiac dysfunction. Collagen content and organization was assessed through picrosirius red staining and fractal dimension (FD) analysis, respectively. Transcript abundance was estimated through reverse transcription-quantitative PCR (RT-qPCR). Cardiac dysfunction was confirmed by the reduction in maximum pulmonary artery velocity and pulmonary artery acceleration time. Through histomorphometric assessment, we found no differences in the interstitial collagen FD between groups. Regarding gene expression, *myh7* gene expression was upregulated in the TPAH group. However, this did not occur with the S group. PAH also increased the mRNA abundance of *col1a1* in the SPAH and TPAH groups. Moreover, the TPAH group showed a higher abundance of this gene when compared to the S group. With these findings, we concluded that preventive exercise had a positive impact on compensated hypertrophy during pulmonary hypertension. This can be explained in part by the modulation of the extracellular matrix and myosin gene expression in trained rats.

## Introduction

Pulmonary Arterial Hypertension (PAH) is a severe and disabling disease that causes right ventricular (RV) remodeling. This is shown by compensatory hypertrophy and subsequent right ventricular heart failure (HF), the latter being the main prognostic determinant and common cause of death [1]. Alterations in myosin and extracellular matrix-related genes are possible mechanisms involved in the PAH cardiac hypertrophy phase. A study of the HF phase in isolated right ventricular myocytes using monocrotaline demonstrated a reduction in ATPase activity in the myosin head [2]. These changes in myosin heavy chains are critical in the different forms of HF. The chains are the main contractile proteins of the heart, and alterations can directly lead to decreased myocardial contractility [2, 3].

Cardiac collagen increases have been shown in other studies that used monocrotaline for induced right ventricular HF [4]. Cardiac collagen increases have been associated with different forms of overload pressure and increases to myosin with lower ATPase capacity [3]. Right ventricular failure is characterized by extensive fibrosis and changes to extracellular matrix protein expression, collagen, and metalloproteinases [5]. However, the genetic expressions of myosins, collagen, and metalloproteinases have not been studied in the compensatory hypertrophy phase of PAH [5].

Exercise is a commonly-used approach to control and limit cardiac damage. It promotes changes in cardiac remodeling and shows benefits in human and animal models with RV hypertrophy [6,7]. Preventive exercise promotes a cardioprotective effect on PAH, as it improves RV function and softens the evolution of the pathological cardiac remodeling process [8,9]. Various molecular mechanisms have been studied to evaluate cardiac functional improvement from preventive training. These mechanisms include the expression of calcium transit genes, regulation of TNF superfamily cytokines, and the quantification of myosin isoforms. However, the effects of changes to the extracellular matrix gene expression and myosins on PAH have not been explored. In the HF phase, pathological remodeling is impossible to reverse by therapy. Thus, approaches to treat compensatory hypertrophy are important to alleviate dysfunctional impairment [10].

PAH-compensated right ventricular hypertrophy often evolves to HF and results in high death rates and frequent hospitalizations. This validates the necessity of elucidating effects of preventive training and molecular mechanisms on right ventricular hypertrophy, as this phase precedes HF. This study hypothesizes that preventive aerobic training mitigates the gene changes in the compensated ventricular hypertrophy phase in monocrotaline-induced PAH rats. The study investigates the influence of preventive aerobic training in rats with compensated right ventricular hypertrophy on the gene expression of myosin heavy chains and the extracellular matrix.

## Materials and methods

### Ethical approval

The experimental protocols used in this study were approved by the Animal Experimentation Ethics Committee (CEUA) from the University of Western São Paulo, Presidente Prudente, São Paulo, Brazil (Protocol numbers 2483 and 2484). The rats received care following the “Laboratory Animal Care Principles” formulated by the National Society for Medical Research and the Guide for the Care and Use of Laboratory Animals, prepared by the Laboratory Animal Research Institute [11].

### Experimental groups

To conduct this study 32 male Wistar rats were used, 2 months of age and average weight of 205±17.43 g, from the Central Animal Facility of the University of Western São Paulo, UNOESTE. All animals were housed in a room under temperature control at 23 °C, relative humidity of 50–60%, and kept on a 12-hour light/dark cycle. Food and water were supplied *ad libitum*.

The animals were randomly distributed into four experimental groups of eight animals each, denominated as Sedentary Control (S; n=8); Control with Training (T; n=8); Sedentary with Pulmonary Arterial Hypertension (SPAH; n=8); and Pulmonary Arterial Hypertension with training (TPAH; n=8).

### Preventive Training

Rats from the T and TPAH groups were submitted to an adapted treadmill aerobic training protocol (model TK 1, IMBRAMED). The protocol consisted of 13 total weeks, five days a week. The first two weeks were for adaptation (pre-training). After, the rats performed the exercises for eight weeks, with gradual increases in intensity, as described previously. The rats were then injected with monocrotaline or saline and performed the exercises for three more weeks. [12, 13].

### Incremental exercise test

The rats in the T and TPAH groups were submitted to incremental stress tests. These were performed 24 hours after monocrotaline administration and at the start of the 11th, 12th, and 13th weeks to adjust the exercise speed [13, 14]. All exercise was performed with 0% slope. The tests began with a warm-up at 0.5 km/h, followed by five minutes of rest. The speed was then increased to 0.7 km/h for three minutes, followed by increases of 0.2 km/h every three minutes until lactate reached a 1 mmol/l comparative value or exhaustion [15]. Exhaustion was defined as the moment when rats could not continue running for three consecutive minutes. After each increased load, the rats were manually removed from the exercise area for one minute for blood collection. Blood samples were taken from rat tails every three minutes. We used an Accutrend Plus lactometer (Roche, Barcelona, Spain). The device was calibrated to the manufacturer’s specifications. The calculation for stipulating maximum velocity was performed using the arithmetic mean of all experimental group velocities until lactate threshold or exhaustion [16]. Lactate threshold was defined as the rate of rotation without a lactate increase of 1.0 mmol/l above the blood-lactate concentration [12,17]. We used an adapted version of the protocol created by Carvalho *et al.* [15].

### Echocardiographic evaluation

Echocardiographic evaluation was performed using an echocardiogram (General Electric Medical Systems, Vivid S6, Tirat Carme, Israel) equipped with a 5-11.5 MHz multifrequency probe. The rats were intraperitoneally anesthetized with ketamine (50mg kg − 1; Dopalen®) and xylazine (0.5 mg kg − 1; Anasedan®).

The following LV variables were measured: diastolic (LVDD) and systolic (LVSD) diameters, ratio of E and A waves (E/A), percentage of endocardial shortening (EFS), isovolumetric relaxation time (IVRT), heart rate frequency (HR), ejection fraction (EF), and posterior wall shortening velocity (EPVP). The following RV variables were measured: pulmonary artery flow obtained by doppler, maximum flow velocity time [Acceleration time velocity (PVAT)], pulmonary ejection time (PET), and peak flow velocity (PVF) [12, 18, 19]. Pulmonary velocity acceleration time is an indicator of the severity of pulmonary hypertension. Increases to pulmonary systolic blood pressure levels correspond to decreases in PVAT values. Pulmonary ejection time is a parameter related to systolic function and the degree of PAH. Maximum flow velocity is related to RV systolic function [12, 20, 21].

### Euthanasia

After the echocardiographic evaluation (48 hours), the rats were weighed and then euthanized with an intraperitoneal dose of sodium pentobarbital (50 mg/kg). At euthanasia two observers determined the presence or absence of clinical and pathological congestive heart failure features. The clinical finding suggestive of heart failure was tachypnea/labored respiration. Pathologic assessment of cardiac decompensation included subjective evaluation of pleuropericardial effusion, atrial thrombi, ascites, and liver congestion.

### Evaluation of anatomical parameters

The heart was removed, dissected into the atria (AT), right ventricles (RV) and left ventricles (LV) and ventricular septum and weighed. The anatomical parameters were normalized by the final body weight (AT/FBW, RV/FBW and LV/ FBW) and were used as the hypertrophy index. The lungs and liver were also removed, weighed and stored in an oven for 48 h. Next, they were weighed again to calculate the wet/dry weight ratio which was used to evaluate signs of cardiac failure [15].

### Histology and fractal analysis

The right ventricle was divided into two parts. One part was fixed in 10% buffered formalin solution for 48 hours, and the other was used for gene expression analysis. After fixation, the tissues were placed on paraffin blocks. Coronal histological sections were viewed using a Leica microtome (RM 2155). The histological sections were stained on slides with haematoxylin–eosin solution (HE) to measure the cross-sectional areas of the cardiomyocytes, using a LEICA microscope (model DM750, Leica Microsystems, Wetzlar, Germany). At least 50 cardiomyocyte diameters were measured from each RV as the shortest distance between borders drawn across the nucleus.

Histological sections of the RV myocardial interstitium were stained on histological slides by the Picrosirius technique for collagen visualization. The cardiac tissue images were captured by a computer coupled to a camcorder. Digital images from the LEICA DM LS microscope (model DM750, Leica Microsystems, Wetzlar, Germany) were sent to a computer equipped with Image-Pro Plus (Media Cybernetics, Silver Spring, U.S.). The red collagen color (picrosirius) were turned blue to reveal the percentage of collagen in relation to the total area of the image. Twenty fields of each right ventricle were analyzed using a 40X objective with 400x magnification. The chosen fields were far from the perivascular region [22].

Binarized photographs and the box-counting method using ImageJ software were used for FD analysis. The software used box-counting with two dimensions. This allowed for the quantification of pixel distribution without interference from the texture of the image. This results in two images (binarized and gray level) with the same FD. The analysis of the fractal histological slides was based on the relation between the resolution and the evaluated scale. The result was quantitatively expressed as the FD of the object with DF ¼ (Log Nr/Log r_1; Nr as the quantity of equal elements needed to fill the original object with scale applied to the object). FD was calculated using the ImageJ software set between 0 and 2, without distinguishing different textures [23, 24, 25].

### Real-time polymerase chain reaction after reverse transcription (RT-qPCR)

Total RNA was extracted from RV tissue using TRIzol (ThermoScientific, Waltham, U.S.) and then treated with DNAse deoxyribonuclease I (ThermoScientific) following the manufacturer’s instructions. RNA integrity was evaluated by agarose gel electrophoresis for visualization of ribosomal RNAs. The High Capacity Reverse Transcriptional Kit (ThermoScientific) was used for the synthesis of complementary RNA (cDNA) from 1000 ng of total RNA for each sample. Using real-time quantitative PCR (qPCR), cDNA was used to evaluate the relative levels of Rattus norvegicus myosin heavy chain 7 (*myh6*) mRNA, Rattus norvegicus myosin heavy chain 7 (*myh7*), Rattus norvegicus myosin heavy chain 7B (*myh7b*), Rattus norvegicus collagen type I alpha 1 chain (*col1a1*) mRNA, Rattus norvegicus collagen type I alpha 2 chain (*col1a2*), Rattus norvegicus collagen type 3 alpha 1 chain (*col3a1*) mRNA, and Rattus norvegicus metalloproteinases 2 (*mmp2*) mRNA. The Taqman™ Universal Master Mix II (AppliedBiosystems, Foster City, U.S.) and the StepOne Plus system (ThermoScientific) were used for qPCR. All samples were analyzed in duplicates. The cycling conditions were at 50 °C for two minutes and 95 °C for 10 minutes. This was followed by 40 cycles of denaturation at 95 °C for 15 seconds and the final extension at 60 °C for one minute. Gene expression was quantified relative to the values of the S group after normalization by expression levels of the beta-actin reference gene (*Actb*) using the 2 ^ -DDCt method [26]. Primer sequences were selected from GenBank transcript access numbers (http://pubmed.com) and designed using Primer Express v.3.0 software (ThermoScientific).

### Data analysis

Statistical analyses were performed using GraphPad Prism software (Graph-Pad Software, La Jolla, U.S.). The Shapiro-Wilk test was used to assess data normality. To analyze the echocardiogram data, Kruskal-Wallis test and Dunn’s post test were used with data from the collagen interstitial fraction and the expression of myh6 and *col1a1* mRNA. ANOVA and Tukey’s post test were used with data from the expression of myh7 and *col1a2* mRNA. Data were expressed with box plot graphs that showed the first and third quartile, median, minimum, and maximum. The significance level was considered when p <0.05.

### Results Echocardiographic evaluation

The LV echocardiographic evaluation is presented in Table 1. LVDD was lower in the SPAH group when compared to rats in the control group. LVDD was higher in the T group when compared to rats in the control group. The rats treated with monocrotaline presented RV dysfunction due to the decrease in the maximum pulmonary artery velocity (Vmax-Pulm) and the pulmonary artery acceleration time (TAC-pulm). An improvement of VMmax-Pulm in the TPAH group was recorded (Figure 1).

**Table 1.** Left ventricle echocardiographic evaluation.

| PARAMETERS | S (n=8) | SPAH (n=8) | T (n=8) | TPAH (n=8) |
| --- | --- | --- | --- | --- |
| HR (bpm) | 310.71±46.58 | 331.66±35.65 | 314±38.25 | 325.44±42.81 |
| LVDD (mm) | 7.84±0.54 | 7.02±0.73* | 8.25± 0.50* | 7.37±0.94 |
| LVSD (mm) | 4.14±0.48 | 3.59±0.47 | 4.39± 0.39 | 3.45±0.81 |
| EFS | 47.33±3.56 | 48.72± 6.05 | 46.86± 2.28 | 53.50± 7.07 |
| E/A | 1.55± 0.31 | 1.07±0.36 | 1.37± 0.19 | 1.12± 0.37 |
| PWSV (mm/s) | 34.21±0.90 | 37.37± 3.43 | 37.85± 3.30 | 37.38±3.85 |
| IVRT (ms) | 22.57± 3.59 | 31.55± 9.40 | 22± 3.57 | 28.66±7.26 |
| Ejection Fraction | 0.85±0.030 | 0.86±0.048 | 0.84±0.019 | 0.89±0.043 |

Data are expressed as mean ± standard deviation. Heart rate (HR), diastolic (LVDD) and systolic (LVSD) diameters, percentage of endocardial shortening (EFS), the ratio of E and A waves (E/A), posterior wall shortening velocity (PWSV), time isovolumetric relaxation rate (IVRT), ejection fraction (EF). S (n=8): Sedentary Control; T (n=8): Control with Training; SPAH (n=8): Sedentary Pulmonary Arterial Hypertension; TPAH (n=8): Pulmonary Arterial Hypertension with Training.
\* Statistical difference $p < 0.05$ .

**Figure 1.**
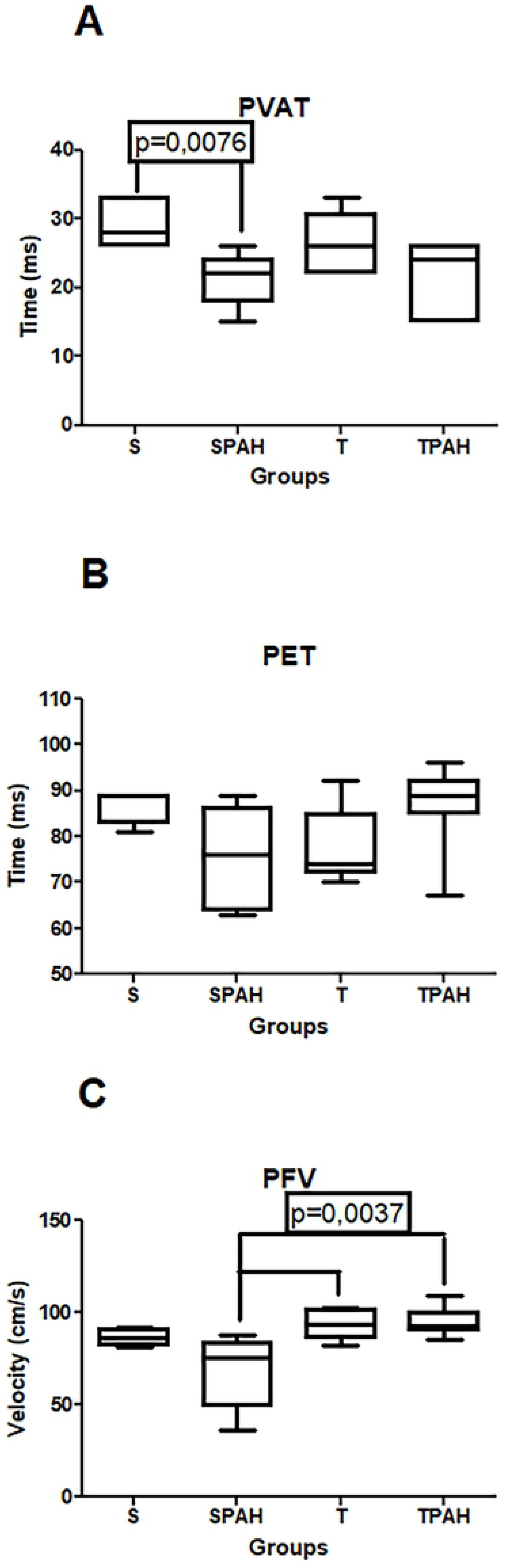
RV echocardiographic evaluation presented in a boxplot. A. PVAT: pulmonary velocity acceleration time; B. PET: pulmonary artery ejection time; C. PVF: peak flow velocity of the pulmonary artery. S (n=8): Sedentary Control; T (n=8): Control with Training; SPAH (n=8): Sedentary Pulmonary Arterial Hypertension; TPAH (n=8): Pulmonary Arterial Hypertension with Training.

### Group characterization and anatomic parameters

In PAH (n=16), all rats presented right ventricular and atria hypertrophy (S= 0,22 ± 0,03 g; SPAH= 0,37 ± 0,18 g; TPAH= 0,36 ± 0,19 g, p < 0,05). There was no clinical or pathological evidence of heart failure.

### Histological and fractal analysis

Fiber cross sectional areas were higher in PAH groups (S= 62,39 ± 6,37 μm^2^; SPAH: 104,88 ± 21,83 μm^2^; TPAH= 89,23 ± 7,99 μm^2^, p < 0,05).Comparisons between rats that performed preventive exercise and sedentary rats did not show statistical difference (p> 0.05) in the percentage of cardiac interstitial collagen (Figure 2) and fractal analysis (Figure 3).

**Figure 2.**
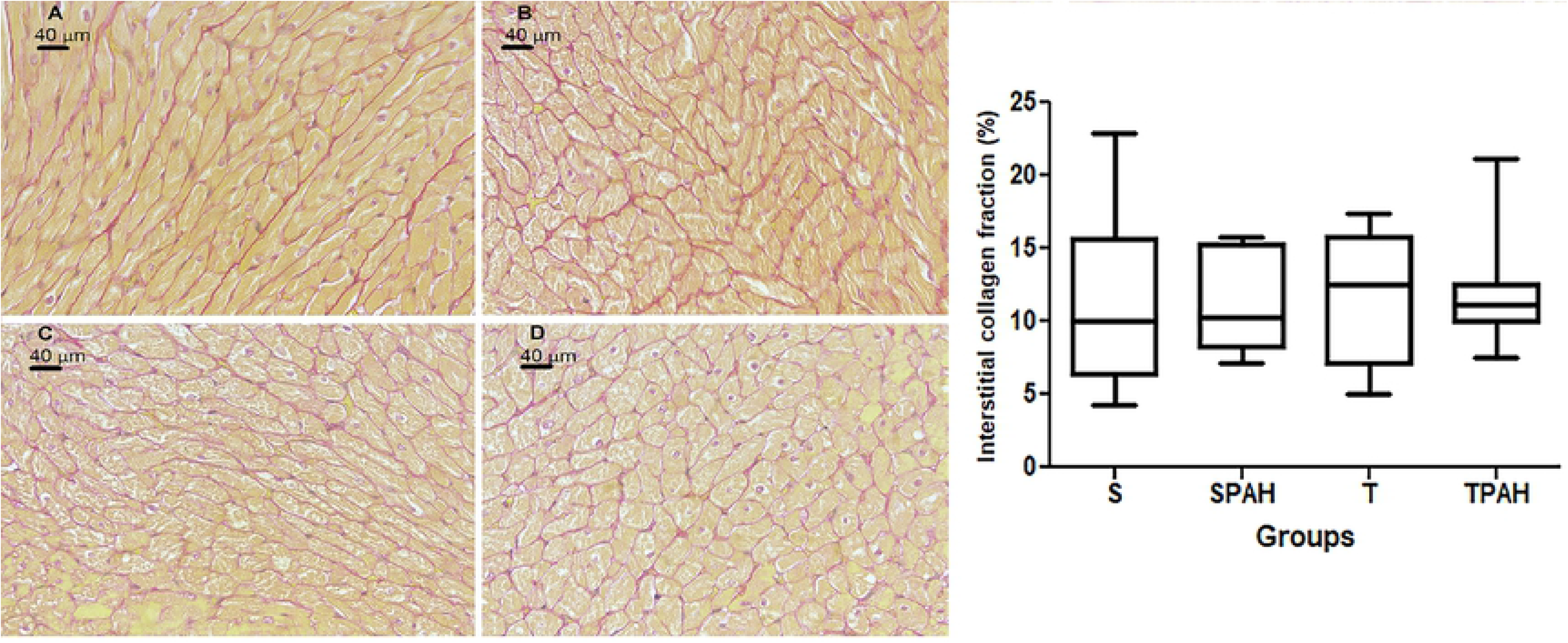
Absence of impact of preventive exercise on cardiac interstitial collagen fraction in PAH. Cross-sections of the cardiac muscle were stained by the Picrosirius Red technique and viewed with 40x objective and 400x magnification. Groups A. S (n=8): Sedentary Control; B. T (n=8): Control with Training; SPAH (n=8): Sedentary Pulmonary Arterial Hypertension; TPAH (n=8): Pulmonary Arterial Hypertension with Training.

**Figure 3.**
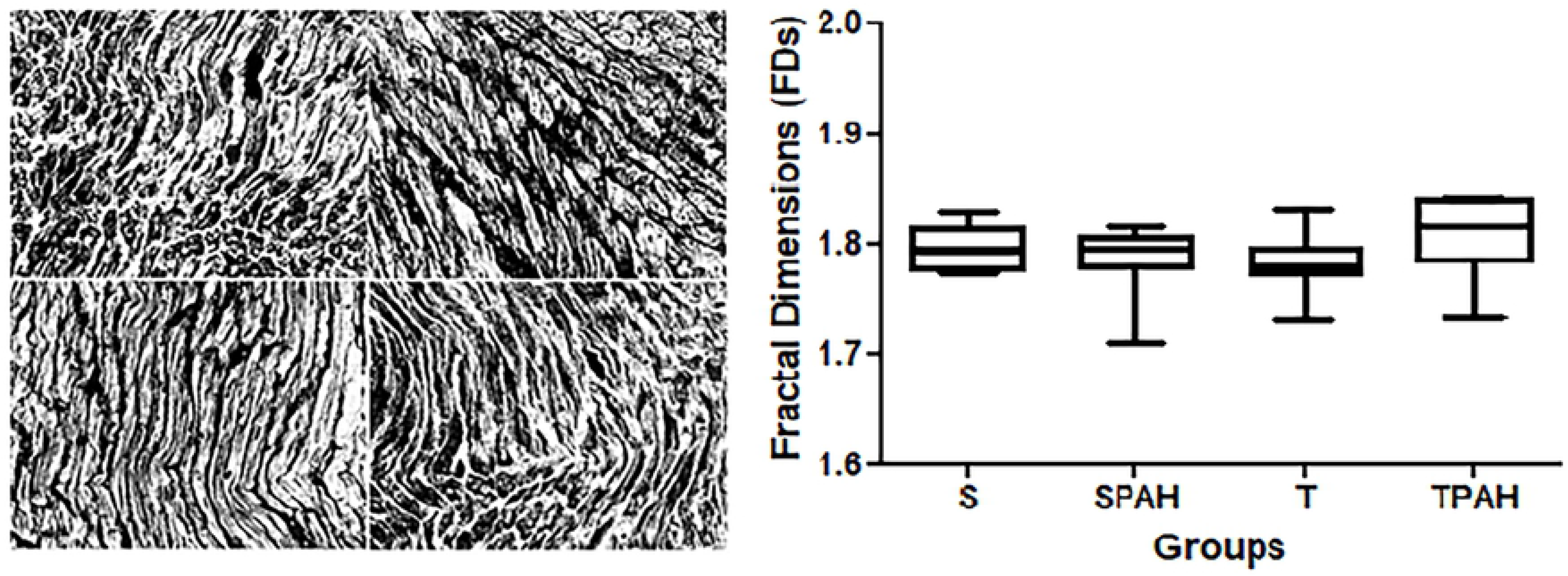
Fractal dimension analysis of right ventricle tissue stained with the Picrosirius Red technique and viewed with 40x objective and 400x magnification.

### Relative gene expression

Preventive exercise increased the Myh7 expression gene in the TPAH group when compared to the control group (S vs. TPAH, p = 0.0242). The expression of *col1a1* was higher in the groups with PAH when compared to the sedentary and trained groups (S vs. SPAH, S vs. TPAH, T vs. TPAH, p = 0.0008). The other groups did not present statistically significant differences (Figure 4).

**Figure 4.**
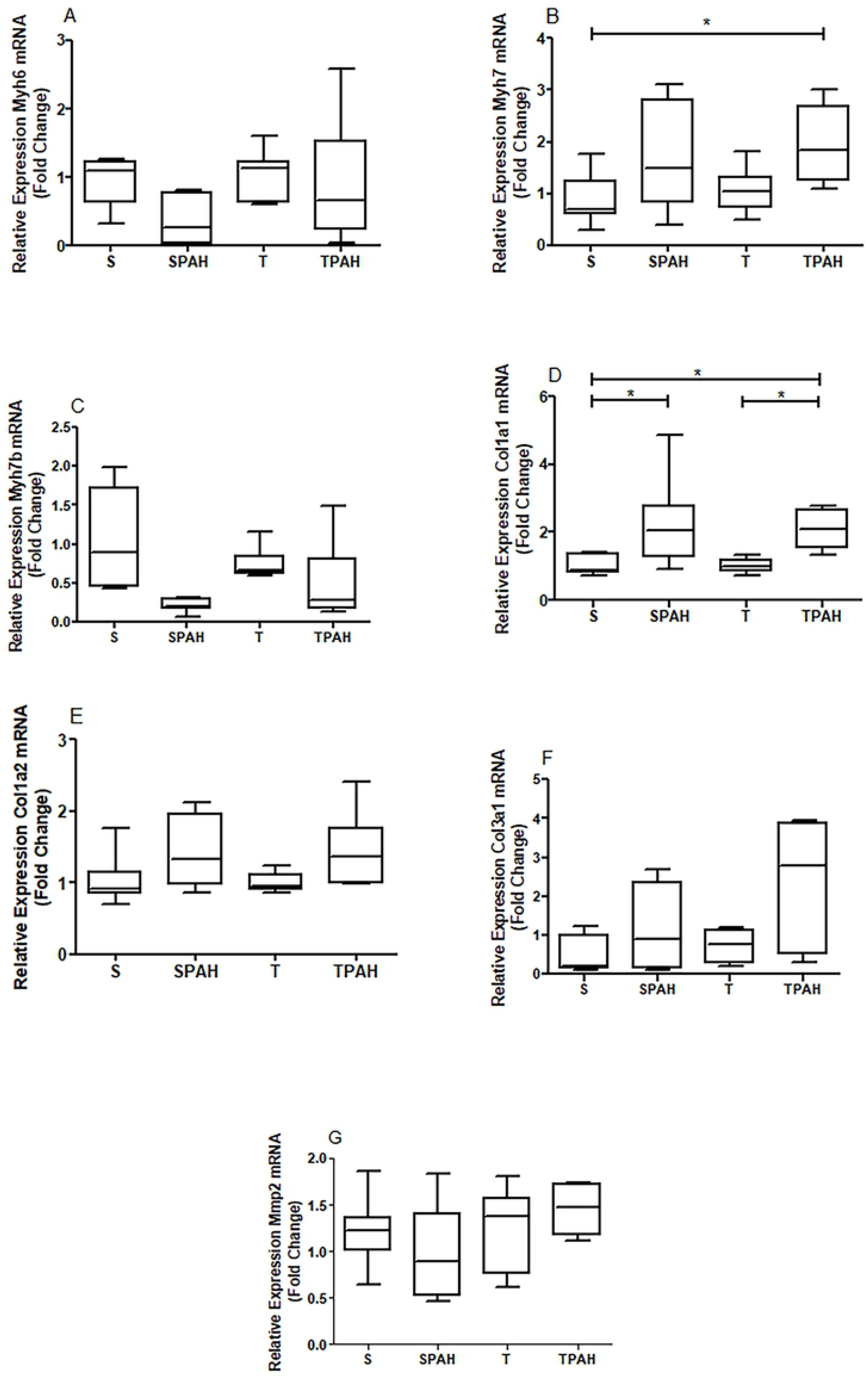
The mRNA abundance of A. *Myh6* gene expression. B. *Myh7* gene expression. C. *Myh7b* gene expression. D. *Col1a1* gene expression. E. *Col1a2* gene expression. F. *Col3a1* gene expression. G. *Mmp2* gene expression in right ventricle of experimental groups. S (n=8): Sedentary Control; T (n=8): Control with Training; SHAP (8): Sedentary Pulmonary Arterial Hypertension; TPAH (n=8): Pulmonary Arterial Hypertension with Training. * Statistical difference p <0.05.

## Discussion

The main finding of the current study was that preventive physical exercise increased *myh7* and *col1a* in the PAH-compensated ventricular hypertrophy phase. This interfered in disease progression. Despite persistent right pressure overload, echocardiography confirmed an increase in cardiac function.

Myosins are the major contractile proteins of muscle. In the heart, there are heavy myosin chains (α-MHC and β-MHC), regulatory light myosin chains, and essential light chain protein C-linked myosin [23, 27]. Α-MHC has a higher sliding speed and two to three times higher ATPase activity than β-MHC; but β-MHC can generate force with lower energy expenditure [28]. Our results showed an increase in the myh7 gene responsible for the β-MHC protein in rats from the PAH preventive exercise group.

The increase in the myh7 gene may be a compensatory mechanism in the PAH exercise group. The increase of this specific myosin is typically related to poorly adaptive cardiac remodeling [29]. However, our study showed that this increase came with cardiac function improvements. A study from Moreira-Gonçalves *et al*. 2015 [4] also showed an increase of myosin in an exercise-performing group that was without weakened ventricular function. Another study showed that rats with HF (from isoproterenol) preserved cardiac function with increased *myh7* and reduced *myh6* [29].

Early expressed transcription factors are other mechanisms that relate to the regulation of MHC genes. These include GATA transcription factor 4, NK2 homeobox 5 (*Nkx2-5-gene*), MADS-box transcription factor, serum response factor (Srf) (attenuates the expression of *myh6* and *myh7*), myocyte enhancer factor 2 (MEF2) and AT, factors that bind to the Mcat malonyl-CoA-acyl carrier protein transacylase sequence, and forkhead box O1 (Foxo1). Foxo1 can act as an *myh6* repressor through histone deacetylase or the N-CoR nuclear suppressor [30]. A purine-rich negative regulatory (PNR) element is present in the first region of the *myh6* gene. When increased, it reduces the expression of this myosin by 20 to 30 times [31, 32]

In addition to myosins, there is a cardiac collagen network that supports and connects all structures and assists inadequate cardiac function (systolic and diastolic). The network has a fundamental role in resisting pathological deformations, maintaining structural alignment, regulating distensibility, and forcing transmission during cardiac fiber shortening [7, 33, 34]. In our study, the phase of cardiac dysfunction by PAH increased *col1a1* gene expression, demonstrating the role of this gene in the worsening of cardiac functionality.

Collagen content and organization were unaltered in our study. FD is a useful method for assessing the organization via images from fractals. These images reveal the amount of space and self-similarity of the structure, detect subtle morphological changes, and perform functional quantitative measurements [25]. Despite alterations to gene expression, fractal evaluation showed that tissue collagen organization was preserved.

In pathological situations, such as acute myocardial infarction and pressure overload, one study showed that increased interstitial fibrosis directly relates to the worsening of ventricular contractile function [34]. The authors of this study also used echocardiography to demonstrate the worsening of contractile function. Size increases to the septal thickness, LV posterior wall, and chamber were observed and associated with reduced ejection fraction and increased diastolic pressure.

In different forms of HF, the gene expressions of *col1a1, col1a2*, and *col3a1* increase due to pressure overload or infarction. However, quantity depends on the cause of HF [34–37]. With pressure overload, an increase of type 1 collagen leads to cardiac stiffness. This occurs from diastole and systole alterations, loss of control in structure alignment, and the regulation of cardiac distensibility and force transmission [7, 33, 34].

Cardiac collagen is altered by pressure overloads from mechanical stress-activating fibroblasts. This induces an inflammatory process through concomitant increases in the extracellular matrix [38]. In the monocrotaline PAH model, pressure overload is from increased pulmonary vascular resistance, with activation of the NF-κB pathway [8]. Our study shows these changes may cause an early increase of the *col1a1* gene.

In response to an injury, the cardiac extracellular matrix assists in electrical and chemical signals, provides structural support, and facilitates mechanical signals. The matrix has metalloproteinases that play an important role in several cardiac pathologies, including dilated cardiomyopathy, myocardial infarction, and hypertensive cardiac hypertrophy [39]. MMPs 2 and 9 cause damage to cardiomyocytes when increased. This leads to the worsening of the cardiac muscle and HF [39]. We did not find the expected increase in MMP2 in the groups with PAH. This factor may have contributed to cardiac function preservation.

Rats with aortic constriction have demonstrated greater increases in the *col1a1* gene when compared to *col3a1* [35]. In acute myocardial infarction, the *col3a1* gene increases more than *col1a1*. This is a consequence of the cardiac tissue healing process [34]. One study demonstrated that both types of collagen genes can be attenuated by post-transcriptional inhibition from miR-29b microRNA. In addition, aerobic training alters a set of microRNAs associated with improved heart function [39]. For the hypertension group in our study, increases to *collal* gene expression and functional worsening were recorded. This gene continued to increase in the exercise group, and functional improvements were recorded. Other cardioprotective factors released in the myocardium from exercise could have neutralized these adverse aspects of cardiac remodeling. These include the expression of genes that control the transport of calcium, regulate TNF superfamily cytokines, improve oxidative function [4, 8], and have anti-fibrotic effects for inflammatory process reduction [40].

Physical exercise is used as a non-pharmacological approach for PAH. Exercise is performed for cardiopulmonary rehabilitation of the disease. However, exercise as a preventive approach for right ventricular dysfunction has not been thoroughly researched [8, 13]. In our study, *myh7* and *col1a1* genes increased and cardiac function improved (observed by echocardiography) after preventive exercise on a treadmill for up to 60 minutes, five days a week for 13 weeks, at speeds until 1.1 km/h [13]. Additional molecular mechanisms should be studied to demonstrate improvements from exercise in the early phase of PAH.

Beneficial influences of various types of exercise on myosin and cardiac collagen have already been demonstrated. Aerobic exercise induces physiological cardiac hypertrophy from the volume load imposed on the heart without rest periods. This results in increased biosynthesis of contractile components, including increases to fast myosin heavy chains (α-MHC) and decreases to slow isoforms (β - MHC) [41].

Regarding collagen, SOCI *et al*. [42] demonstrated that swimming increases miRNA-29c expression in healthy rats, reducing the expression of cardiac collagen genes *col1a1* and *col3a1*. Another study showed rats with cardiac abnormalities from aging reduced fibrosis and *colla2* after exercise on the treadmill for 12 weeks [43]. For acute myocardial infarction, exercise on the treadmill with a moderate inclination of 5° for 45 minutes reduced *col1a2* and *col3a1* [44]. In rats six weeks after acute myocardial infarction, resistance exercise four times a week with 75% of 1RM with 10-12 repetitions combined with treadmill exercise five times a week at 15 meters/minute for 12 weeks improved the interstitial collagen fraction due to the effect of pathological hypertrophy reversal [45, 46]. However, increased interstitial collagen was not observed in our study. The increase in the *col1a1* gene may indicate that collagen increases later and affects cardiac dysfunction. Similar to myosins, collagen may be influenced by exercise modality, frequency, duration, intensity, and the disease phase when training starts [8].

We conclude from our findings that preventive exercise is beneficial for compensated hypertrophy in pulmonary hypertension. This could be partially explained by the modulation of the extracellular matrix and myosin gene expression in exercise group rats. Other mechanisms, pathways, and variations in exercise intensity and type should be investigated in the preclinical PAH phase.

Lastly, our findings showed that exercise yielded benefits when started before and in the early stages of the disease. This confirmed the importance of exercise as a preventive approach provides considerable cardioprotection against the deleterious effects of PAH, which reinforces the importance of maintaining a physically active lifestyle. Clinical trials that examine effects of exercise on PAH should use individuals with the early stage of the disease.

## Acknowledgments

We would like to thank Eric Schloeffel for his help with English editing.

## References

1 Humbert M, Guignabert C, Bonnet S, et al. Pathology and pathobiology of pulmonary hypertension: state of the art and research perspectives. Eur Respir J. 2019; 53: 1801887

2 Vescovo G, Jones SM, Harding SE & Poole-wilson PA. Isoproterenol sensitivity of isolated cardiac myocytes from rats with monocrotaline-induced right-sided hypertrophy and heart failure. J Mol Cell Cardiol. 1989; 21: 1047–61.

3 Batlle M, Castillo N, Alcarraz A, et al. Axl expression is increased in early stages of left ventricular remodeling in an animal model with pressure-overload. PLoS One. 2019;14(6): e0217926.

4 Moreira-Gonçalves D, Ferreira R, Fonseca H, et al. Cardioprotective effects of early and late aerobic exercise training in experimental pulmonary arterial hypertension. Basic Res Cardiol. 2015; 110:57.

5 Nadadur RD, Umar S, Wong G, et al. Reverse right ventricular structural and extracellular matrix remodeling by estrogen in severe pulmonary hypertension. J Appl Physiol. 2012; 113:149–158.

6 Colombo R, Siqueira R, Becker CU, et al. Effects of exercise on monocrotaline-induced changes in right heart function and pulmonary artery remodeling in rats. Can J Physiol Pharmacol. 2013; 91: 38–44.

7 Zile MR, Baicu CF, Ikonomidis J, et al. Myocardial Stiffness in Patients with Heart Failure and a Preserved Ejection Fraction: Contributions of Collagen and Titin. Circulation. 2015; 131: 1247–1259.

8 Nogueira-Ferreira R, Moreira-Gon D, SilvaAna F, et al. Exercise preconditioning prevents MCT-induced right ventricle remodeling through the regulation of TNF superfamily cytokines, Intern J Cardiol. 2016; 203:858–866.

9 Pacagnelli FL, Aguiar AF, Campos DHS, et al. Training improves the oxidative phenotype of muscle during the transition from cardiac hypertrophy to heart failure without altering MyoD and myogenin. Exp Physiol. 2016; 101:1075–1085.

10 Gonzalez A, Ravassa S, Beaumont J, Lopez B, Diez J. New targets to treat the structural remodeling of the myocardium. J Am Coll Cardiol. 2011; 58:1833–1843.

11 Clark J.D., Gebhart G.F., Gonder J.C. et al. (1997) The 1996 guide for the care and use of laboratory animals. ILAR J. 38, 41–48.

12 Lopes FS, Carvalho RF, Campos GE, Sugizaki MM, Padovani CR, Nogueira CR, Cicogna AC, Dal-Pai-Silva M. Down-regulation of MyoD gene expression in rat diaphragm muscle with heart failure. Int J Exp Pathol. 2008; 89: 216–222.

13 Pacagnelli FL, Sabela AK, Okoshi K, et al. Preventive physical training exerts a cardioprotective effect in rats treated with monocrotaline. Int j Exp Pathol. 2016; 97: 238–247.

14 Rodrigues B, Figueroa DM, Mostarda CT, Heeren MV, Irigoyen MC & Angelis K. Maximal exercise test is a useful method for physical capacity and oxygen consumption determination in streptozotocin-diabetic rats. Cardiovasc diabetol. 2007; 13: 1–7.

15 Carvalho JF, Masuda MO & Pompeu FAMS. Method for diagnosis and control of aerobic training in rats based on lactate threshold. Comp Biochem Physiol A Mol Integr Physiol. 2005; 140:409–413.

16 Svedah K & Macintosh BR. Anaerobic threshold: the concept and methods of measurement. Canad J Appl Physiol. 2003; 28: 299–323.

17 Souza RWA, Piedade WP, Soares LC, et al. Aerobic exercise training prevents heart failure-induced skeletal muscle atrophy by anti-catabolic, but not anabolic actions. PLoS One. 2014; 9: 1–15.

18 Ferreira JCB, Rolim NPL, Bartholomeu JB, Gobatto CA, Kokubun E & Brum PC. Maximal lactate steady state in running mice: effect of exercise training. Clin Exp Pharmacol Physiol. 2007; 34: 760–765.

19 Martinez ST, Santos APB & Pinto AC. A Determinação Estrutural do Alcaloide Pirrolizidínico Monocrotalina: Exemplo dos Desafios da Química de Produtos Naturais Até os Anos Sessenta do Século XX. Rev Virtual Quim. 2013; 5: 300–311.

20 Eguchi M, Ikeda S, Kusumoto S et al. Adipose-derived regenerative cell therapy inhibits the progression of monocrotaline-induced pulmonary hypertension in rats. Life Sci. 2014; 118: 306–312.

21 Dabestani A, Mahan G, Gardin JM et al. Evaluation of pulmonary artery pressure and resistance by pulsed Doppler echocardiography. Am J Cardiol. 1987; 59:662–668.

22 Pacagnelli FL, Okoshi K, Campos DHS, et al. Physical training attenuates cardiac remodeling in rats with supra-aortic stenosis. Exp Clin Cardiol. 2014; 20:3889–3905.

23 Zornoff LAM, Matsubara BB, Matsubara LS & Minicucci MF. Cigarette smoke exposure intensifies ventricular remodeling process following myocardial infarction. Arq Bras Cardiol. 2006; 86:276–282.

24 Cury SS, Freire PP, Martinucci B, et al. Fractal dimension analysis reveals skeletal muscle disorganization in mdx mice. Biochem Biophys Res Commun. 2018; 3; 503 (1):109–115.

25 Fávero PF, Lima VAV, Santos PH, et al. Differential fractal dimension is associated with extracellular matrix remodeling in developing bovine corpus luteum. Biochem Biophys Res Commun. 2019; 27; 516(3): 888–893.

26 Livak KJ & Schmittgen KD. Analysis of relative gene expression data using real-time quantitative PCR and the 2 (DeltaDeltaC(T)) method. Methods. 2001; 25, 402–408.

27 Marsiglia JDC & Pereira AC. Cardiomiopatia Hipertrófica: Como as Mutações Levam à Doença? Arq Bras Cardiol. 2014; 102, 295–304.

28 Cai M, Huang Q, Liao W, Wu Z, Liu F & Gao Y. Hypoxic training increases metabolic enzyme activity and composition of myosin heavy chain isoform in rat ventricular myocardium. Eur J Appl Physiol. 2010; 108:105–111.

29 Kralova E, Doka G, Pivackova L, Srankova J, et al. L-Arginine Attenuates Cardiac Dysfunction, But FurtherDown-Regulates a-Myosin Heavy Chain Expression inIsoproterenol-Induced Cardiomyopathy. Basic & Clinic Pharmacol Toxicol. 2015; 117:251–260

30 Qi Y, Zhu Q, Zhang K, et al. Activation of Foxo1 by Insulin Resistance Promotes Cardiac Dysfunction and β–Myosin Heavy Chain Gene Expression. Circ Heart Fail. 2015; 8: 198–208.

31 Gupta MP. Factors controlling cardiac myosin-isoform shift during hypertrophy and heart failure. J Mol Cell Cardiol. 2007; 43: 388–403.

32 Giger JM, Bodell PW, Baldwin KM, Haddad F. The CAAT-binding transcription factor 1/nuclear factor 1 binding site is important in β-myosin heavy chain antisense promoter regulation in rats. Exp Physiol. 2009; 94:1163–1173.

33 Shoulders MD & Raines RT. Collagen structure and stability. Annu Rev Biochem. 2009; 78: 929–958.

34 Zornoff LAM, Paiva SAR, Duarte DR & Spadaro J. Remodelação Ventricular Pós-Infarto do Miocárdio: Conceitos e Implicações Clínicas. Arq Bras Cardiol. 2009; 92: 157–164.

35 Yang F, Li P, Li H, Shi Q, Li S & Zhao L. microRNA-29b Mediates the Antifibrotic Effect of Tanshinone IIA in Postinfarct Cardiac Remodeling. J Cardiovasc Pharmacol. 2015; 65: 456–64.

36 Wang JH, Su F, Wang S, et al. CXCR6 deficiency attenuates pressure overload-induced monocytes migration and cardiac fibrosis through downregulating TNF-α-dependent MMP9 pathway. Int J Clin Exp Pathol. 2014; 7: 6514–23.

37 Wang Q, Yu X, Xu H, Zhao X, Sui D, Re G. Improves Isoproterenol-Induced Myocardial Fibrosis and Heart Failure in Rats, Evidence-Based Complem and Alternat Med. 2019; 1–9.

38 Lindner D, Zietsch C, Tank J, et al. Cardiac fibroblasts support cardiac inflammation in heart failure. Basic Res Cardiol. 2014; 109:428.

39 Souza RWA, Fernandez GJ, Cunha JPQ, et al. Regulation of cardiac microRNAs induced by aerobic exercise training during heart failure. Am J Physiol Heart CircPhysiol. 2015; 309: 1629–1641.

40 Agarwal D, Haque M, Sriramula S, Mariappan N, Pariaut R, Francis J. Role of proinflammatory cytokines and redox homeostasis in exercise-induced delayed progression of hypertension in spontaneously hypertensive rats. Hypertension. 2009; 54:1393–1400.

41 Mcmullen JR & Jennings GL. Differences between pathological and physiological cardiac hypertrophy: novel therapeutic strategies to treat heart failure. Clin Exp Pharmacol Physiol. 2007; 34:255–262.

42 Soci UP, Fernandes T & Hashimoto NY. MicroRNAs 29 are involved in the improvement of ventricular compliance promoted by aerobic exercise training in rats. Physiol Genomics. 2011; 43: 665–73.

43 Kwak HB, Kim JH, Joshi K, Yeh A, Martinez DAn & Lawler JM. Exercise training reduces fibrosis and matrix metalloproteinase dysregulation in the aging rat heart. Faseb J. 2011; 25: 1106–17.

44 Xu X & Wan W. Effects of exercise training on cardiac function and myocardial remodeling in post myocardial infarction rats. J Mol Cell Cardiol. 2008; 44: 114–22.

45 Nunes RB, Alves JP, Kessler LP & Dal Lago P. Aerobic exercise improves the inflammatory profile correlated with cardiac remodeling and function in chronic heart failure rats. Clinics. 2013; 68: 876–82.

46 Alves JP, Nunes RB, Stefani GP & Dal Lago P. Resistance training improves hemodynamic function, collagen deposition and inflammatory profiles: experimental model of heart failure. PLoS One. 2014; 9:e110317.

